# NLR immune receptor-nanobody fusions confer plant disease resistance

**DOI:** 10.1101/2021.10.24.465418

**Authors:** Jiorgos Kourelis, Clemence Marchal, Sophien Kamoun

**Affiliations:** The Sainsbury Laboratory, University of East Anglia, Norwich Research Park, NR4 7UH, Norwich, UK

**Author notes:** These authors contributed equally to this work (Jiorgos Kourelis, Clemence Marchal).

**Keywords:** NLR, NB-LRR, NBS-LRR, NLR-ID, Innate immunity, Adaptive immunity

## Abstract

Plant pathogens cause recurrent epidemics that threaten crop yield and global food security. Efforts to retool the plant immune system have been limited to modifying natural components and can be nullified by the emergence of new pathogen races. Therefore, there is a need to develop made-to-order synthetic plant immune receptors with resistance tailored to the pathogen genotypes present in the field. Here we show that plant immune receptors can be used as scaffolds for VHH nanobody fusions that bind fluorescent proteins (FPs). The receptor-nanobody fusions signal in the presence of the corresponding FP and confer resistance against plant viruses expressing FPs. Given that nanobodies can be raised against virtually any molecule, immune receptor-nanobody fusions have the potential to generate resistance against all major plant pathogens and pests.

**One-Sentence Summary:** Plant immune receptor-nanobody fusions enable made-to-order disease resistance genes.

## INTRODUCTION

Plants lack an adaptive immune system an rely on innate immune receptors to detect invading pathogens. Therefore, efforts to retool the plant immune system to design new-to-nature biochemical activities have been relatively limited to modifying natural components, for instance through receptor mutagenesis or domain shuffling (Kim *et al*., 2016; Pottinger *et al*., 2020; Segretin *et al*., 2014; Giannakopoulou *et al*., 2015; De la Concepcion *et al*., 2019; Cesari *et al*., 2021; Wang *et al*., 2021; Dangl *et al*., 2013). While these approaches have yielded promising results, they often target a specific pathogen isolate and thus lack plasticity and adaptability to a wider range of pathogens and pests. In addition, plant pathogens are notorious for rapidly evolving virulent races that can nullify new resistance specificities. Thus, there is a need for an adaptative system where new resistance can be engineered as required to target the pathogen genotypes associated with plant disease outbreaks.

One class of immune proteins that could be optimal templates for receptor engineering are the subset of intracellular NOD-like/nucleotide-binding leucine-rich repeat immune receptors (NLRs) that carry unconventional integrated domains (IDs) (Cesari *et al*., 2014; Wu *et al*., 2015; Kroj *et al*., 2016; Sarris *et al*., 2016). These IDs are generally thought to mediate pathogen effector detection, either by directly binding to effectors or by acting as a substrate for their enzymatic activity; and this activity is subsequently translated into an immune response (Cesari *et al*., 2013; Sarris *et al*., 2015; Le Roux *et al*., 2015; Maqbool *et al*., 2015; Guo *et al*., 2018). Often these NLR-containing IDs (NLR-IDs) are genetically linked to conventional NLRs that are required for immune activation following effector detection (Eitas and Dangl, 2010; Cesari *et al*., 2014). Pik-1 and Pik-2 are such an NLR receptor pair from rice carrying an N-terminal coiled-coil (CC) domain (Ashikawa *et al*., 2008). Pik-1 carries an integrated heavy metal associated (HMA) domain between its CC and the central NB-ARC (nucleotide-binding domain shared with APAF-1, various R-proteins and CED-4) domains that directly binds AVR-Pik effector proteins secreted by the blast fungus *Magnaporthe oryzae* (Kanzaki *et al*., 2012; Cesari *et al*., 2013; Maqbool *et al*., 2015; De la Concepcion *et al*., 2018; De la Concepcion, Maidment, *et al*., 2021; Białas *et al*., 2021; Białas *et al*., 2017). Activated Pik-1 relies on its partner Pik-2, a MADA-type CC-NLR, which is thought to function through oligomerization into a resistosome structure similar to the Arabidopsis NLR ZAR1 (Adachi *et al*., 2019; Wang, Wang, *et al*., 2019; Wang, Hu, *et al*., 2019). The integrated HMA domain of Pik-1 can be mutated to confer new pathogen effector responses, however, this has only been described for effectors from the blast fungus (De la Concepcion *et al*., 2019).

What would be the ultimate integrated domain for engineering made-to-order plant immune receptors? Given that animal adaptive immunity has the capacity to generate antibodies against virtually any antigen it is exposed to, we reasoned that harnessing antibodies for plant immunity would potentially enable building receptors that respond to any plant pathogen molecule. We focused on the minimal antigen-binding fragment of single-domain heavy chain antibodies (known as VHHs or nanobodies) of camelid mammals (Hamers-Casterman *et al*., 1993; Greenberg *et al*., 1995; Muyldermans, 2013; Könning *et al*., 2017), because they are small soluble 10-15 kDa domains, that tend to correctly fold intracellularly and have many useful properties for biotechnological applications. To test our idea, we generated orthogonal Pik-1 sensors in which the integrated HMA domain is swapped with nanobodies that bind either GFP or mCherry (Kirchhofer *et al*., 2010; Fridy *et al*., 2014) **(Fig. 1A, Table S1)**. We hypothesized that the engineered versions of Pik-1 would trigger immunity in the presence of GFP or mCherry.

**Fig. 1.**
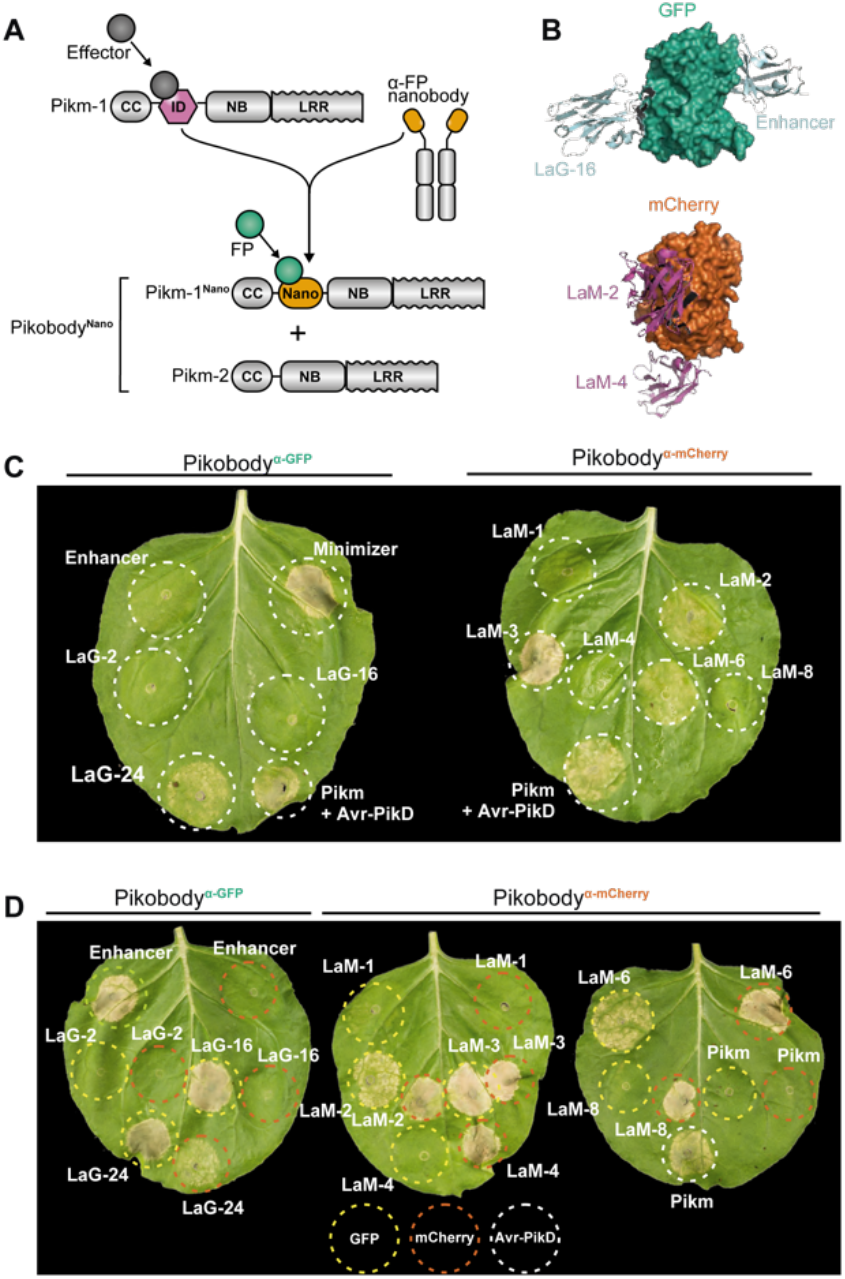
NLR immune receptor-nanobody fusions trigger a hypersensitive cell death response in presence of the corresponding fluorescent protein antigen. (**A**) Engineering of fluorescent protein (FP)-activated NLR sensor. The integrated HMA domain of the NLR Pikm-1, which is involved in pathogen effector recognition by direct binding, was swapped with nanobodies binding GFP of mCherry. (**B**) Structures of the interaction interfaces of GFP (green) and the GFP-binding nanobodies Enhancer (3K1K) (Kirchhofer *et al*., 2010) and LaG-16 (6LR7) (Zhang *et al*., 2020) (light green) or mCherry (orange) and the mCherry-binding nanobodies LaM-2 (6IR2) and LaM-4 (6IR1) (purple). (**C**) Screen for autoimmunity of engineered Pikobody^α- GFP/α-mCherry^ upon overexpression in *N. benthamiana*. Representative *N. benthamiana* leaves infiltrated with appropriate constructs photographed 5 days after infiltration. The infiltration site for each construct is labelled on the picture and circled with a white dashed line. Autoimmunity is characterized by a hypersensitive cell death response (HR, cell-death) at the infiltration site (e.g. Minimizer), whereas no autoimmunity is observed if the leaf remains green at the infiltration site (e.g. Enhancer). Pikm-1/Pikm-2 pair (Pikm on the figure) co-infiltrated with AVR-PikD was used as a positive control for HR. (**D**) Pikobody^α-GFP/α-mCherry^ co-expressed with the corresponding fluorescent protein antigen results in a specific HR. Representative *N. benthamiana* leaves infiltrated with appropriate constructs photographed 5 days after infiltration. The infiltration site for each construct is labelled on the picture and circled with a yellow, orange or white dashed line where GFP, mCherry or AVR-PikD was co-infiltrated with the Pikobody, respectively. Pikm-1/Pikm-2 pair (Pikm on the figure) co-infiltrated with AVR-PikD was used as a positive control for HR.

## RESULTS

Engineered NLR-IDs often exhibit autoimmune activities in the absence of a ligand (Białas *et al*., 2021; De la Concepcion, Benjumea, *et al*., 2021). This is possibly due to structural changes induced when new domains or point mutations perturb the resting state of the receptor. Hence, we first tested whether the Pik-1-nanobody fusions induce autoimmunity. Of the 11 tested Pik-1-nanobody fusions, six didn’t exhibit autoimmunity when expressed with Pik-2 in leaves of the model plant *Nicotiana benthamiana* (**Fig. 1B**), indicating that they can be used for follow-up gain-of-function assays. Next, we co-expressed these six fusions with GFP and mCherry. Among these, four produced a hypersensitive cell death response (HR, immune response readout) specifically when expressed with their matching fluorescent proteins (FPs) (**Fig. 1C**, Enhancer, LaG-16, LaM-4, and LaM-8). Additionally, a further three fusions which displayed weak autoimmunity gave a stronger hypersensitive cell death only when combined with their matching fluorescent proteins (**Fig. 1B, C**, LaG-24, LaM-2, and LaM-6). This indicates that the Pik-1-VHH fusions are functional and can be endowed with new-to-nature activities. We termed this engineered immune receptor system Pikobody (**Fig. 1**).

We selected Pikobody^Enhancer^ (Pikm-2/Pikm-1^Enhancer^) and Pikobody^LaM4^ (Pikm-2/Pikm-1^LaM4^), recognising GFP and mCherry, respectively, to further confirm our results. We first challenged the absence of autoimmunity of these two Pikobodies by co-expressing them in *N. benthamiana* leaves with the gene silencing inhibitor p19, which is known to elevate heterologous expression levels (van der Hoorn *et al*., 2003) (**Fig. S1**). Whereas Pikobody^LaM4^ did not produce any detectable HR, we could score a weak response in 5 out of 18 samples for Pikobody^Enhancer^ when co-expressed with p19 (**Fig. S1**). These results confirm that these two Pikobodies are mainly inactive on their own and are appropriate for further experiments. Next, we co-expressed Pikobody^Enhancer^ and Pikobody^LaM4^ with GFP and mCherry and determined that they produced a hypersensitive cell death only with their matching FP and at levels similar to the response obtained with a natural combination of Pikm and a blast fungus effector (**Fig. S2, Fig. S3**). We also noted that Pikobody^Enhancer^ and Pikobody^LaM4^ specifically responded to three pathogen effectors only when they were tagged with the matching GFP/EGFP or mCherry/mRFP1 (**Fig. S4, Fig. S5, Fig. S6**). This further confirmed that the Pikobodies are functional FP sensors that detect FPs even when they are fused to other proteins.

Can Pikobodies produce a functional immune response that is effective against a pathogen? We used recombinant *Potato virus X* (PVX) (Marillonnet *et al*., 2008) expressing either GFP or mCherry to assay the ability of Pikobodies to reduce viral load (**Table S1**). These PVX variants express FPs from a duplicated coat protein sub-promoter in the virus genome. We used fluorescence intensity and immunodetection of GFP/mCherry accumulation as proxy for viral load in leaf samples (**Fig. 2**). Both Pikobody^Enhancer^ and Pikobody^LaM4^ specifically reduced fluorescent intensity of PVX expressed GFP or mCherry, respectively, to an extent comparable to that of Rx, an NLR known to confer immunity against PVX (Bendahmane *et al*., 1999) (**Fig. 2A, B**). This reduction of fluorescence intensity correlates with reduced accumulation of virus expressed GFP or mCherry as compared to the empty vector control or wild-type Pikm-1/Pikm-2 (**Fig. 2C, D**). We did, however, observe a faint signal corresponding to GFP or mCherry in the samples with PVX-GFP or PVX-mCherry and Pikobody^Enhancer^ or Pikobody^LaM4^, respectively, as compared to no detectable FP bands in the samples with Rx (**Fig. 2C, D**). In response to the matching PVX-FP, both Pikobodies triggered a clearly visible hypersensitive cell death further confirming they can produce a potent immune response to the recombinant viruses (**Fig. 2E, F**).

**Fig. 2.**
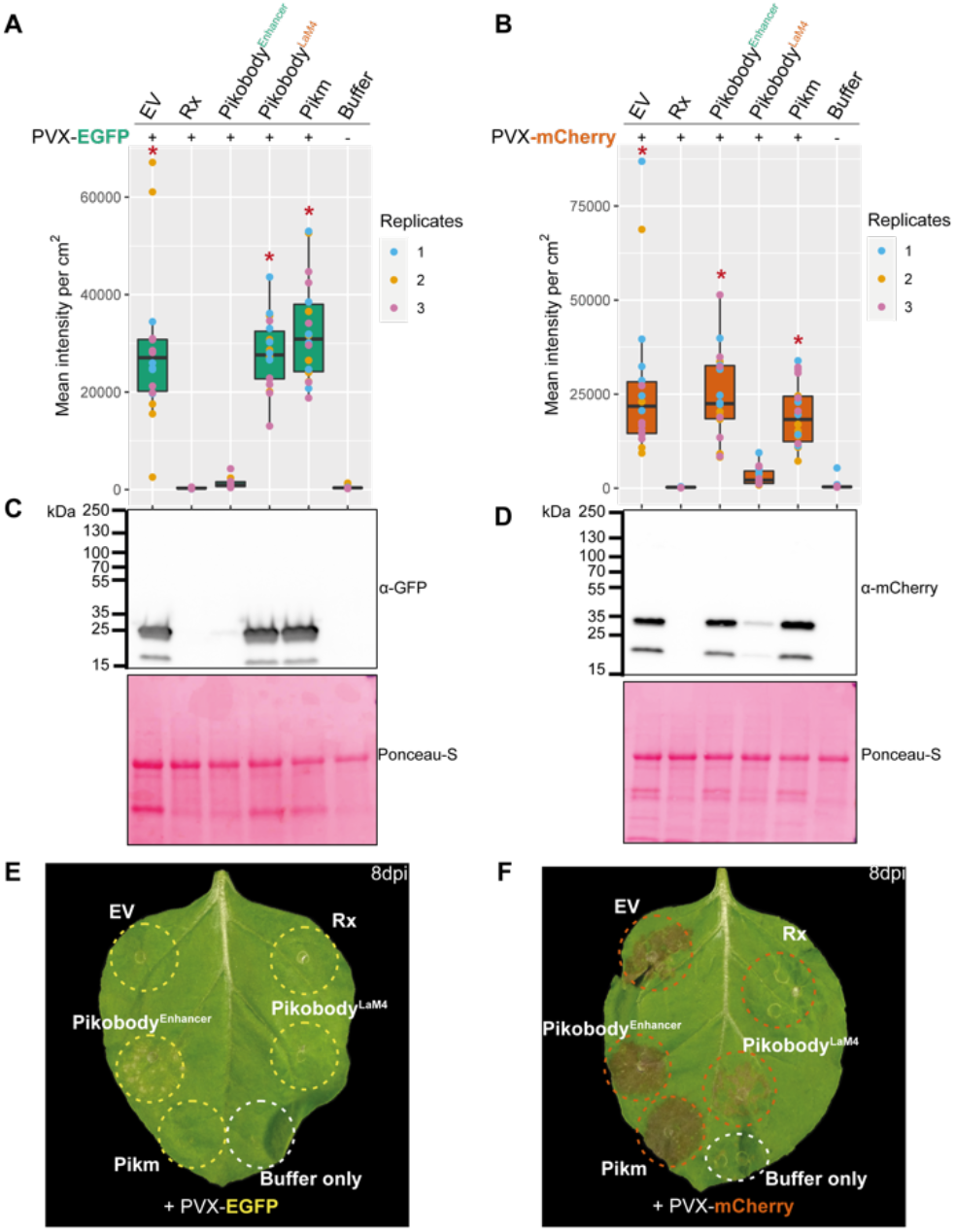
Pikobodies confer resistance against engineered Potato virus X strains expressing matching fluorescent proteins. (**A**-**B**) Specific reduction in fluorescence intensity of PVX expressed EGFP (GFP) or mCherry in the presence of Pikobody^Enhancer^ or Pikobody^LaM-4^, respectively. GFP (**A**) or mCherry (**B**) mean fluorescence intensity per cm^2^ measured in *N. benthamiana* leaves 4 days post-infiltration and used as a proxy for PVX viral load in each infiltration site. The boxplot summarizes data obtained in three independent experiments (replicates) with six infiltration site per construct and per experiment. Red asterisks show significant differences between buffer only (no PVX added, negative control) and tested constructs in the presence of PVX-GFP (**A**) or PVX-mCherry (**B**) (Dunnett’s test, pvalue < 0.001). Empty vector was used as a positive control for viral load and buffer only as a negative control for viral load. Rx, a resistance gene recognizing the coat protein from PVX, was used as a positive control for PVX resistance. (**C-D**) Specific reduction of PVX expressed GFP or mCherry accumulation in the presence of Pikobody^Enhancer^ or Pikobody^LaM- 4^, respectively. GFP or mCherry accumulation was used as proxy to evaluate viral load from PVX-GFP (**C**) or PVX-mCherry (**D**). For the immunoblot analysis, total protein was extracted 4 days after inoculation with PVX-GFP (**C**) or PVX-mCherry (**D**) in the presence of the tested constructs. Empty vector was used as a positive control for viral load and Buffer only as a negative control. GFP and mCherry were detected using anti-GFP or anti-mCherry antibody, respectively. Ponceau S staining shows equal protein loading across samples. (**E-F**) Specific HR triggered by PVX expressed GFP or mCherry in the presence of Pikobody^Enhancer^ or Pikobody^LaM- 4^, respectively. Representative *N. benthamiana* leaves infiltrated with appropriate constructs photographed 8 days after infiltration. The infiltration site for each construct is labelled on the picture and circled with a yellow, red or white dashed line for PVX-GFP (**E**), PVX-mCherry (**F**) or no PVX, respectively. The purple color on the infiltration sites in (**F**) is due to a high accumulation of mCherry.

To independently confirm these results, we tested two additional recombinant PVX variants spliced in different ways to express GFP (Lu *et al*., 2003; Cruz *et al*., 1996) (**Table S1, Fig. S7**). Pikobody^Enhancer^, but not Pikobody^LaM4^, markedly reduced GFP fluorescence intensity and protein accumulation when challenged with a PVX with the GFP sequence inserted between the triple gene block and coat protein in the virus genome (**Fig. S7A-B**). Pikobody^Enhancer^ didn’t significantly affect fluorescence intensity when challenged with a PVX carrying a GFP in-frame fusion to the N-terminus of the virus coat protein and with an inserted porcine teschovirus-1 2A self-cleaving peptide (**Fig. S7C**). However, Pikobody^Enhancer^, but not Pikobody^LaM4^ and other control treatments, reduced the accumulation of virus expressed GFP (**Fig. S7D**). This discrepancy could be explained by the observation that a GFP fraction remains fused to coat protein, therefore somehow affecting fluorescence (**Fig. S7D**) (Cruz *et al*., 1996). Nonetheless, Pikobody^Enhancer^, but not Pikobody^LaM4^, caused a visible hypersensitive cell death with both PVX-GFP (**Fig. S7E, F**). We conclude that Pikobody^Enhancer^ can provide enhanced resistance against multiple PVX-GFP variants.

The addition of more than one immune receptor in a plant variety—a plant breeding strategy known as R gene stacking— can maximize resistance durability in the field by delaying the emergence of virulent pathogen races (Luo *et al*., 2021; Zhu *et al*., 2012; Ghislain *et al*., 2019). However, co-expression of plant immune receptors can lead to autoimmunity (Chae *et al*., 2014; Tran *et al*., 2017) or suppression of recognition (Hurni *et al*., 2014). We investigated whether Pikobodies with different FP specificities are compatible with each other **(Fig. 3)**.We first determined that none of three mis-matched combinations of Pikobodies induced autoimmunity **(Fig. S8**). Co-expression of Pikobody^Enhancer^ or Pikobody^LaM4^ with the wild-type Pikm pair triggered hypersensitive cell death only in presence of GFP or mCherry, respectively, whereas co-expression of Pikobody^Enhancer^ and Pikobody^LaM4^ produced hypersensitive cell death in the presence of both GFP and mCherry **(Fig. 3A-B**, **Fig. S9A**). Similarly, co-expression of Pikobody^Enhancer^ and Pikobody^LaM4^ markedly reduced fluorescence intensity and protein levels of both GFP and mCherry produced by PVX-FPs (**Fig. 3C-F**). The combination of Pikobody^Enhancer^ and Pikobody^LaM4^ also resulted in hypersensitive cell death in response to either PVX-GFP or PVX-mCherry (**Fig. S9B**). We conclude that Pikobody stacking can expand the recognition and response profile of these immune receptors.

**Fig. 3.**
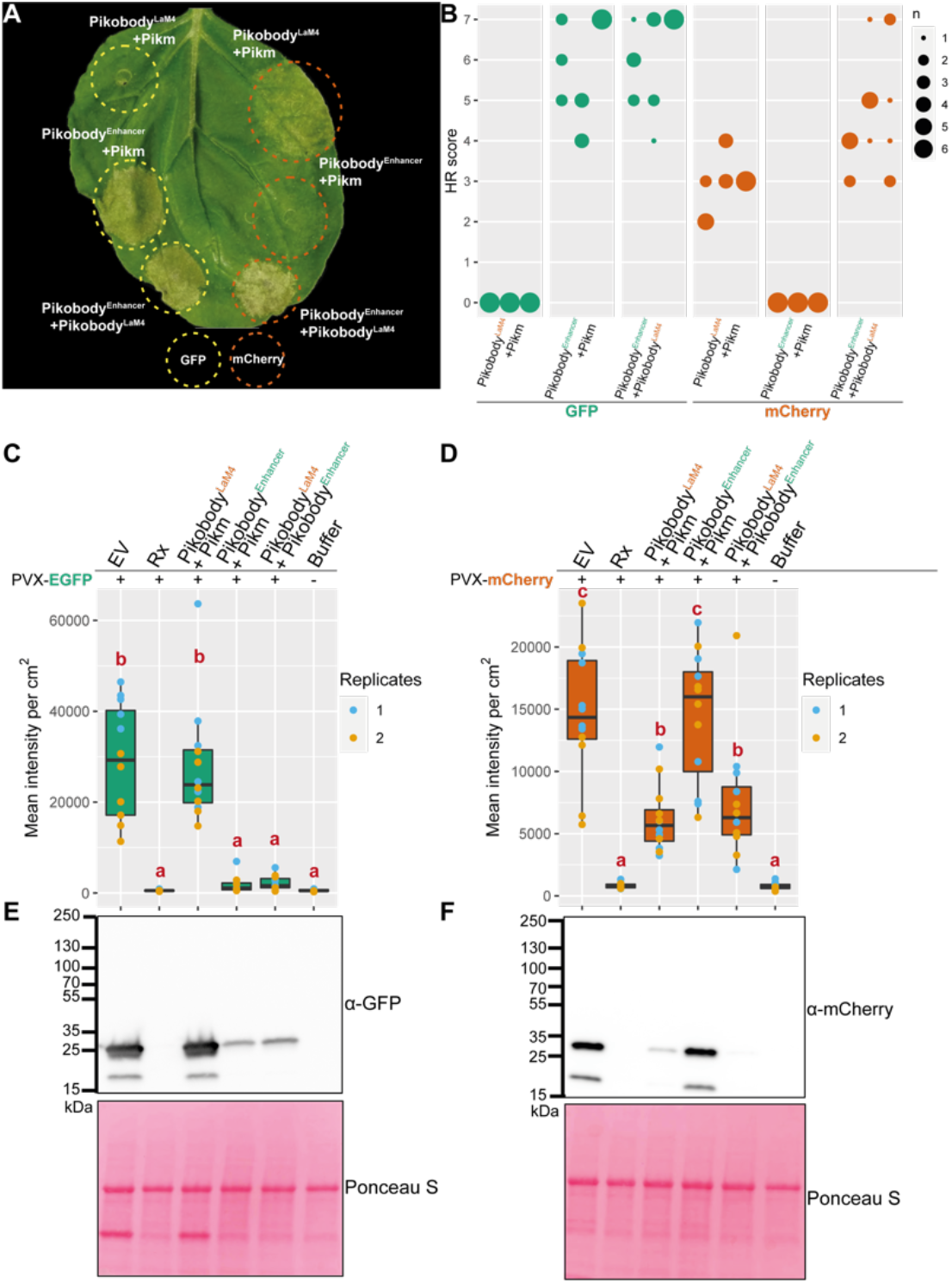
Pikobodies are suitable for stacking. (**A**) Pikobody stacking results in additive immune recognition. Representative *N. benthamiana* leaf infiltrated with constructs indicated in white and infiltration sites circled with a yellow or orange dashed line for GFP or mCherry, respectively. Leaves were photographed 4 days after infiltration. (**B**) HR scores visualized as dots plot, where the size of a dot is proportional to the number of samples with the same score (n) within the same replicate (1 to 3). The experiment was repeated three times with six internal replicates; each column represents a combination of constructs (labelled on the bottom). The statistical analyses of these results are presented in **Fig. S8** and the negative controls are shown in **Fig. S9**. (**C-D**) Specific reduction in fluorescence intensity of PVX expressed GFP and mCherry in the presence of stacked Pikobody^Enhancer^ and Pikobody^LaM-4^. GFP (**C**) or mCherry (**D**) mean fluorescence intensity per cm^2^ measured in *N. benthamiana* leaves 4 days post-infiltration used as a proxy for PVX viral load in each infiltration site. The boxplot summarizes data obtained in two independent experiments (replicates) with six infiltration site per construct and per experiment. Empty vector was used as a positive control for viral load and buffer only as a negative control for viral load. Rx, a resistance gene recognizing the coat protein from PVX, was used as a positive control for PVX resistance. We performed two statistical tests on the presented data: a) Dunnett’s test (pvalue < 0.001) showed significant differences between buffer only (no PVX added, negative control) and Empty vector, Pikobody^LaM-4^+ Pikm in the presence of PVX-GFP (C) and showed significant differences between buffer only and all tested constructs but Rx in the presence of PVX-mCherry (**C**); b) ANOVA followed by Tukey HSD (pvalue < 0.05, letters depict the different groups) showed significant differences between buffer only (no PVX added, negative control) and Empty vector, Pikobody^LaM-4^+ Pikm in the presence of PVX-GFP (C) and showed significant differences between buffer only and all tested constructs but Rx in the presence of PVX-mCherry and significant differences between Pikobody^LaM-4^+ Pikm, Pikobody^LaM-4^+ Pikobody^Enhancer^ and Empty vector, Pikobody^Enhancer^+ Pikm (D). (**E-F**) Specific reduction of PVX expressed GFP and mCherry accumulation in the presence of stacked Pikobody^Enhancer^ and Pikobody^LaM-4^. GFP or mCherry accumulation was used as proxy to evaluate viral load from PVX-GFP (**E**) or PVX-mCherry (**F**). For the immunoblot analysis, total protein was extracted 4 days after inoculation with PVX-GFP (**E**) or PVX-mCherry (**F**) in the presence of the tested constructs. Empty vector was used as a positive control for viral load and Buffer only as a negative control. GFP and mCherry were detected using anti-GFP or anti-mCherry antibody, respectively. Ponceau S staining shows equal protein loading across samples.

We investigated the extent to which Pikobodies function through similar mechanisms as the wild-type Pik pair (Zdrzałek *et al*., 2020) and other CC-NLRs. The conserved P-loop motif within the NB-ARC domain of CC-NLRs is required for the ADP/ATP switch that enables oligomerization into resistosome complexes (Wang, Wang, *et al*., 2019; Dinesh-Kumar *et al*., 2000). Pikobody^K217R/Enhancer^ and Pikobody^K217R/LaM4^ with a P-loop dead mutation in Pikm-2 (Pikm-2^K217R^) failed to produce an HR to their corresponding FP even though the Pikm1 and Pikm2 proteins accumulated to similar levels as the wild-type immune receptors (**Fig. S10**). We conclude that the P-loop motif is required for Pikobody activity, and that the Pikobody system probably functions through the established mechanistic model of CC-NLRs.

## CONCLUSION

We built upon our growing understanding of the evolution and function of the Pik pair of NLRs (Białas *et al*., 2021; Zdrzałek *et al*., 2020; De la Concepcion *et al*., 2018; De la Concepcion *et al*., 2019; De la Concepcion, Maidment, *et al*., 2021) to use Pik-1 as a chassis for VHH nanobody fusions to engineer functional disease resistance genes with new-to-nature functionalities. This strategy for engineering synthetic immune receptors contrasts with earlier approaches, which were based on the modification of endogenous sequences and domains. The Pikobody system can be used for providing a pseudo-adaptive immune system to plants. Given that nanobodies can be readily generated to bind virtually any antigen, Pikobodies have the potential to produce made-to-order resistance genes against any pathogen or pest that delivers effectors inside host plant cells (**Fig. 4**).

**Figure 4:**
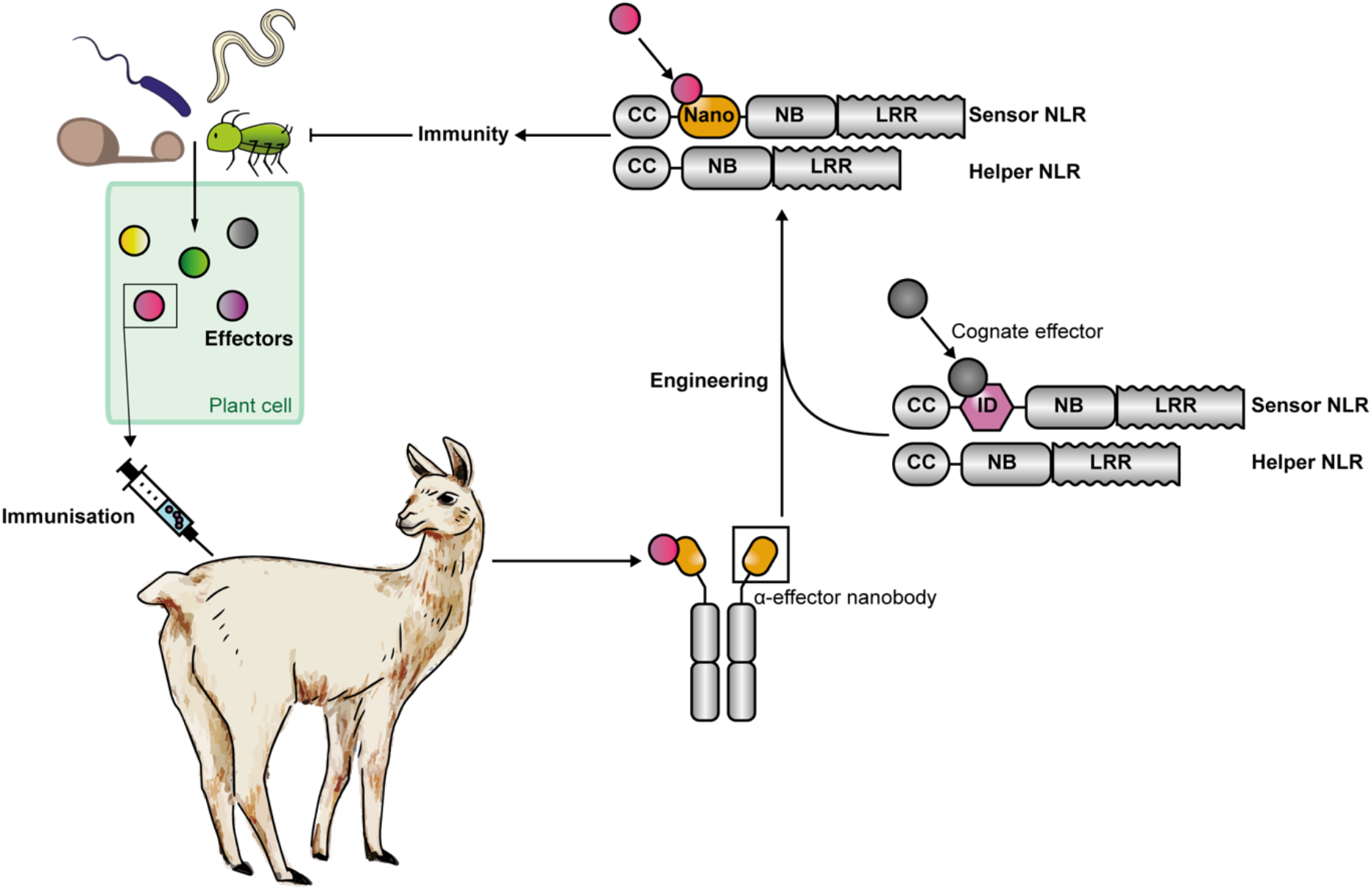
Pipeline for the generation of made-to-order functional Pikobodies. Immunization of camelids with translocated pathogen effectors generates nanobodies which can be integrated into the Pikm scaffold in order to generate functional Pikobodies providing immunity against pathogens translocating these effectors.

## MATERIAL & METHODS

### Plant material and growth conditions

Wild type *N. benthamiana* were grown in Levingtons F2 compost in a glasshouse with set to 24 °C day / 22 °C night, humidity 45–65%, with supplementary lighting when the weather conditions required it.

### Plasmid constructions

The Golden Gate Modular Cloning (MoClo) kit (Weber *et al*., 2011) and the MoClo plant parts kit (Engler *et al*., 2014) were used for cloning, and all vectors are from this kit unless specified otherwise. Cloning design and sequence analysis were done using Geneious Prime (v2021.2.2; https://www.geneious.com). Plasmid construction is described in **Table S1**.

### Transient gene-expression and cell death assays

Transient gene expression in *N. benthamiana* were performed by agroinfiltration according to methods described by van der Hoorn *et al*. (2000). Briefly, *A. tumefaciens* strain GV3101 pMP90 carrying binary vectors were inoculated from glycerol stock in LB supplemented with appropriate antibiotics and grown O/N at 28 °C until saturation. Cells were harvested by centrifugation at 2000 × *g*, RT for 5 min. Cells were washed once and resuspended in infiltration buffer (10 mM MgCl_2_, 10 mM MES-KOH pH 5.6, 200 μM acetosyringone) to the appropriate OD_600_ (see **Table S1**) in the stated combinations and left to incubate in the dark for 2h at RT prior to infiltration into 5-week-old *N. benthamiana* leaves. Two leaves from three plants were inoculated per experiment (N = 6), and the experiment was repeated three times. Hypersensitive cell death phenotypes were scored 4-5 days post-infiltration in a range from 0 (no visible necrosis) to 7 (fully confluent necrosis) according to Adachi *et al*. (2019). The data was visualized with ggplot2 (v3.3.4; Ginestet, 2011) and the statistical analysis was performed using the R package besthr (v0.2.0; MacLean, 2020) (**Table S2** for raw data).

### Protein immunoblotting

Six *N. benthamiana* leaf discs (8 mm diameter) taken 2 days post agroinfiltration were homogenised in extraction buffer [10% glycerol, 25 mM Tris-HCl, pH 7.5, 1 mM EDTA, 150 mM NaCl, 2% (w/v) PVPP, 10 mM DTT, 1x protease inhibitor cocktail (SIGMA), 0.2% IGEPAL® CA-630 (SIGMA)]. The supernatant obtained after centrifugation at 5,600 x g for 10 min at 4 °C was used for SDS-PAGE. 4x SDS-PAGE sample buffer [final concentration: 50 mM Tris-HCl (pH 6.8), 100 mM DTT, 2% SDS, 0.01% bromophenol blue, 10% glycerol] was added and the samples were denatured by incubating at 72 °C for 10 min. Proteins were separated on 10-20%, SDS-PAGE gels (Bio-Rad) and transferred onto polyvinylidene difluoride (PVDF) membrane using a Trans-Blot turbo transfer system (Bio-Rad). Membranes were blocked in in 3% dried milk dissolved in Tris-buffered Saline [50mM Tris-HCL (pH7.5), 150mM NaCl] supplemented with 1% Tween® 20 for 30 min before probing the membrane with either rat monoclonal anti-HA antibody (3F10, Roche) or mouse monoclonal ANTI-FLAG® antibody conjugated to HRP (M2, Sigma) in a 1:4000 dilution. Chemiluminescent detection of signals after addition of either Pierce™ ECL Western (Thermo Fisher Scientific), or 1/5 SuperSignal™ West Femto Maximum Sensitivity Substrate (34095, Thermo Fisher Scientific) was done using the ImageQuant LAS 4000 luminescent imager (GE Healthcare Life Sciences). Equal loading was validated by staining the PVDF membranes with Ponceau S.

### PVX infection assays

Four to five-weeks old *N. benthamiana* plants were inoculated with engineered versions of *Potato virus X* (PVX) (Marillonnet *et al*., 2008; Lu *et al*., 2003; Cruz *et al*., 1996) expressing either GFP or mCherry by agroinfiltration as described above. The PVX expression constructs are detailed in **Table S1**. Two leaves from three plants were inoculated per experiment (N = 6) as in **Fig. 2E, F**, and the experiment was repeated three times. GFP or mCherry fluorescence was measured 4-5 days post-infiltration (see **Table S3** for raw data) using the ImageQuant LAS 4000 luminescent imager (GE Healthcare Life Sciences) with the appropriate excitation and emission settings (EX: epi-RGB, EM:510DF10 filter for GFP and EX: epi-Green, EM: 605DF40 for mCherry). The mean fluorescence intensity per cm^2^ in each infiltration site was measured using Fiji (ImageJ v2.1.0), and a Dunnett’s test was conducted using the R package DescTools (v0.99.41; Signorell, 2021) to compare mean fluorescence intensity per cm^2^ between buffer only (negative control) and all tested constructs followed by an ANOVA and Tukey HSD post-hoc test using the R package agricolae (v1.3-5). The data was visualized using ggplot2 (v3.3.4; Ginestet, 2011).

To determine GFP or mCherry accumulation upon PVX inoculation six leaf discs from the same infiltration site at the same time-point were taken and processed as above. Samples containing PVX expressed GFP or mCherry were diluted threefold in SDS-PAGE sample buffer to prevent saturation of the signal during detection. GFP and mCherry were probed with either mouse monoclonal anti-GFP antibody conjugated to HRP (B-2, Santa Cruz Biotech) or mouse monoclonal anti-mCherry TrueMAB™ antibody conjugated to HRP (OTI10G6, Thermo Fisher Scientific) at a 1:4000 or 1:2500 dilution, respectively. Chemiluminescent detection of signals was done as above.

## ACKNOWLEDGEMENTS

We thank Hsuan Pai for drawing the llama on Figure 4, Aleksandra Białas for helpful comments on the figures, Andrea Williams for useful comments on the text, and Phil Robinson for photography. We also thank our long-standing collaborators Mark Banfield, Nick Talbot and Ryohei Terauchi and other members of the BLASTOFF community for the many useful discussions and suggestions.

## AUTHOR CONTRIBUTIONS

Conceptualization: JK, CM, SK; Data curation: CM, JK; Formal analysis: CM, JK; Investigation: JK, CM; Supervision: SK; Funding acquisition: SK; Project administration: SK; Writing initial draft: CM, JK, SK; Editing: CM, JK, SK.

## SUPPLEMENTAL DATA

**Fig. S1.**
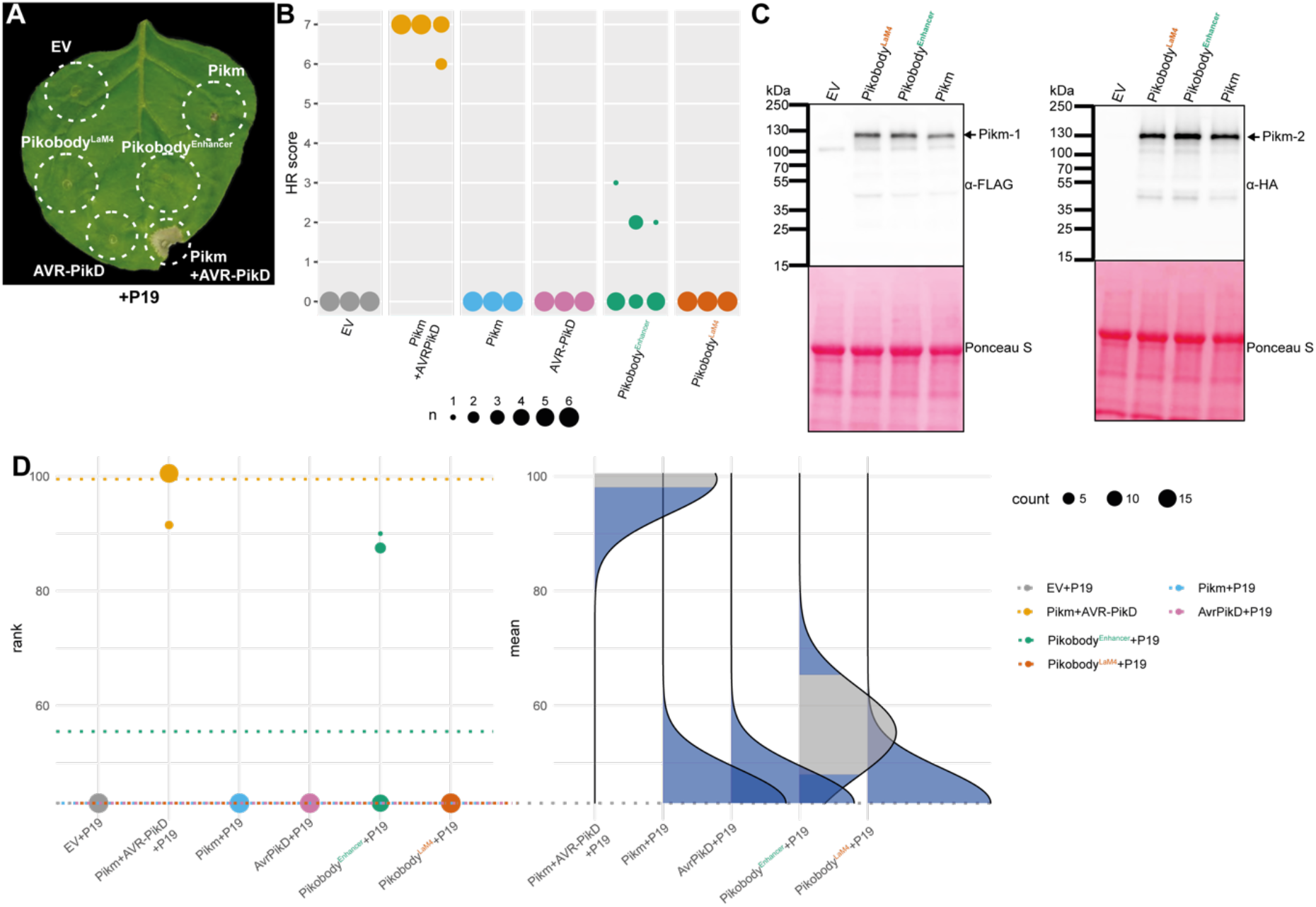
Pikobody^Enhancer^ and Pikobody^LaM4^ are not autoimmune. (**A**) Representative *N. benthamiana* leaf infiltrated with constructs indicated in white and infiltration sites circled with a white dashed line, photographed 4 days after infiltration. (**B**) HR scores visualized as dots plot, where the size of a dot is proportional to the number of samples with the same score (n) within the same replicate (1 to 3). The experiment was repeated three times with six internal replicates; each column represents a combination of constructs (labelled on the bottom). The statistical analyses of these results are presented in (**D**). (**C**) Swapping the HMA domain for nanobodies does not affect protein accumulation. For the immunoblot analysis, total protein was extracted 2 days after transient expression of Pikobody^Enhancer^, Pikobody^LaM4^ and the wild-type Pikm pair. Pikm-1 and the Pikm-1 Enhancer and LaM-4 were C-terminally 6xHis-3xFLAG-tagged and detected using anti-FLAG antibody, while Pikm-2 was C-terminally 6xHA-tagged and detected using anti-HA antibody. Empty vector was used as a negative control. The black arrow points to the band corresponding to Pikm-1 on the left and Pikm-2 on the right. (**D**) Statistical analysis of hypersensitive response showed in (**B**) conducted using an estimation method using besthr R library (MacLean, 2020). Empty vector combined with the silencing inhibitor p19 was used as a negative control, while the wild-type Pikm pair with the cognate effector AVR-PikD and the silencing inhibitor p19 was used as a positive control for HR. The left panels represent the ranked data (dots) and their corresponding mean (dashed line), with the size of a dot proportional to the number of observations with that specific value. The panels on the right show the distribution of 1000 bootstrap sample rank means, with the blue areas illustrating the 0.025 and 0.975 percentiles of the distribution. The difference is considered significant if the ranked mean for a given condition falls within or beyond the blue percentile of the mean distribution for another condition.

**Fig. S2.**
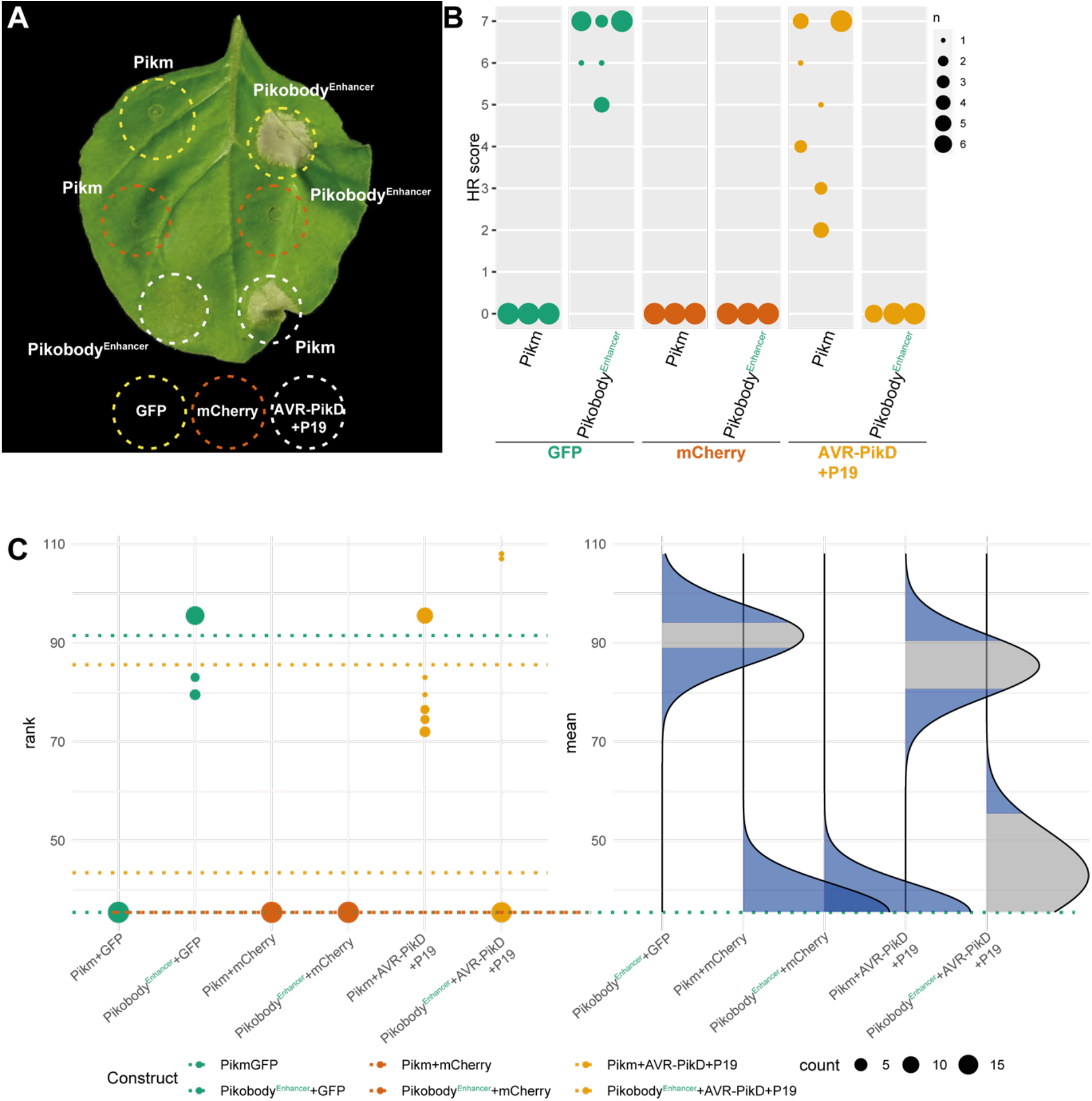
Pikobody^Enhancer^ specifically triggers HR in response to GFP. (**A**) Pikobody^Enhancer^ specifically recognizes GFP. Representative *N. benthamiana* leaf infiltrated with constructs indicated in white and infiltration sites circled with a dashed line, photographed 4 days after infiltration. Yellow, Orange or white color shows sites co-infiltrated with GFP, mCherry or AVR-PikD+P19, respectively. (**B**) HR scores visualized as dots plot, where the size of a dot is proportional to the number of samples with the same score (n) within the same replicate (1 to 3). The experiment was repeated three times with six internal replicates; each column represents a combination of constructs (labelled on the bottom). (**C**) Statistical analysis of hypersensitive response shown in (**B**) conducted using an estimation method using besthr R library (MacLean, 2020). The wild-type Pikm pair with either GFP or mCherry was used as a negative control, while the wild-type Pikm pair with the cognate effector AVR-PikD and the silencing inhibitor p19 was used as a positive control for HR. The left panels represent the ranked data (dots) and their corresponding mean (dashed line), with the size of a dot proportional to the number of observations with that specific value. The panels on the right show the distribution of 1000 bootstrap sample rank means, with the blue areas illustrating the 0.025 and 0.975 percentiles of the distribution. The difference is considered significant if the ranked mean for a given condition falls within or beyond the blue percentile of the mean distribution for another condition.

**Fig. S3.**
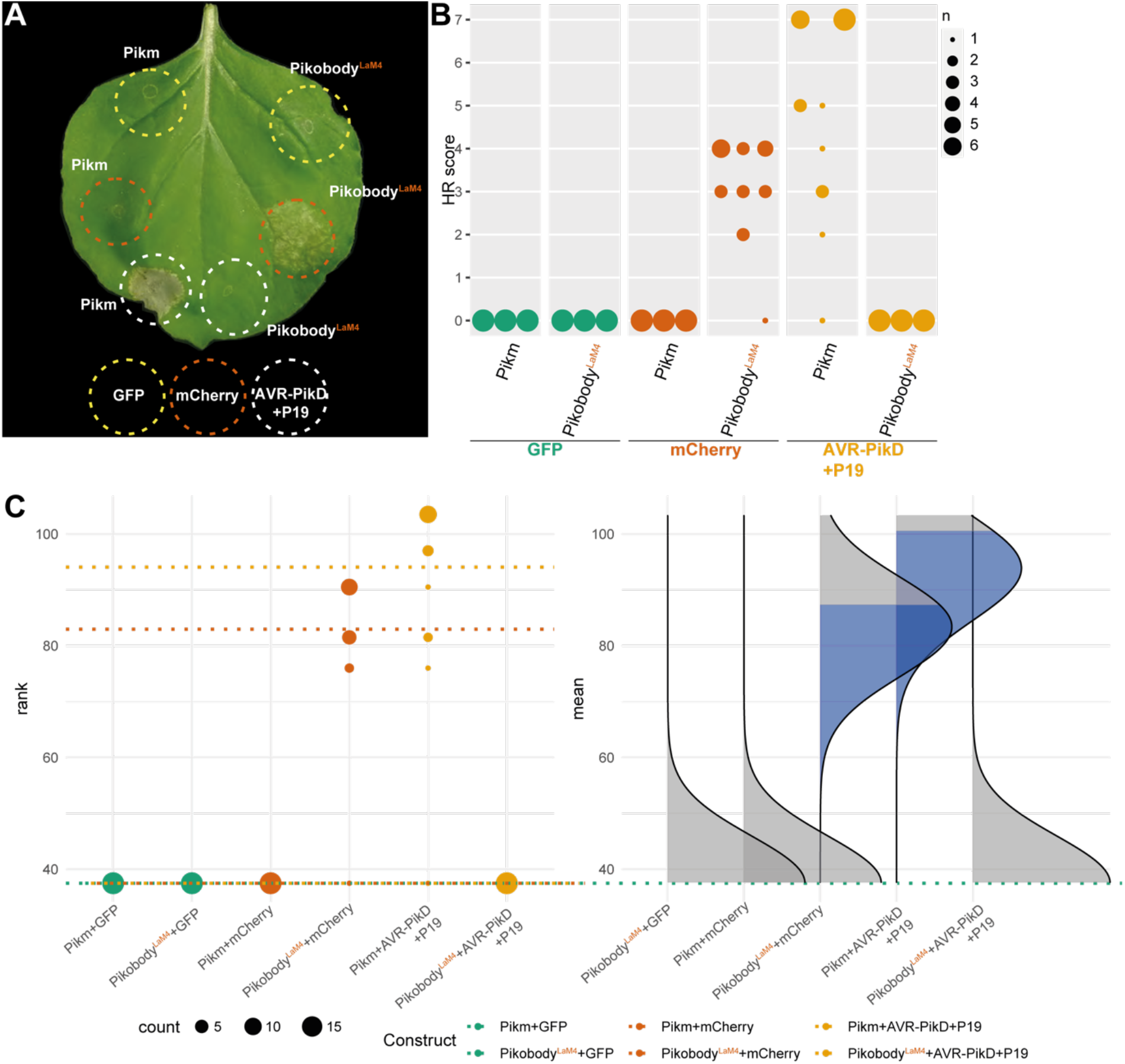
Pikobody^LaM4^ specifically triggers HR in response to mCherry. (**A**) Pikobody^LaM-4^ specifically recognizes mCherry. Representative *N. benthamiana* leaf infiltrated with constructs indicated in white and infiltration sites circled with a dashed line, photographed 4 days after infiltration. Yellow, Orange or white color show sites co-infiltrated with GFP, mCherry or AVR-PikD+P19, respectively (**B**) HR scores visualized as dots plot, where the size of a dot is proportional to the number of samples with the same score (n) within the same replicate (1 to 3). The experiment was repeated three times with six internal replicates; each column represents a combination of constructs (labelled on the bottom). (**C**) Statistical analysis of hypersensitive response shown in (**B**) conducted using an estimation method using besthr R library (MacLean, 2020). The wild-type Pikm pair with either GFP or mCherry was used as a negative control, while the wild-type Pikm pair with the cognate effector AVR-PikD and the silencing inhibitor p19 was used as a positive control for HR. The left panels represent the ranked data (dots) and their corresponding mean (dashed line), with the size of a dot proportional to the number of observations with that specific value. The panels on the right show the distribution of 1000 bootstrap sample rank means, with the blue areas illustrating the 0.025 and 0.975 percentiles of the distribution. The difference is considered significant if the ranked mean for a given condition falls within or beyond the blue percentile of the mean distribution for another condition

**Fig. S4.**
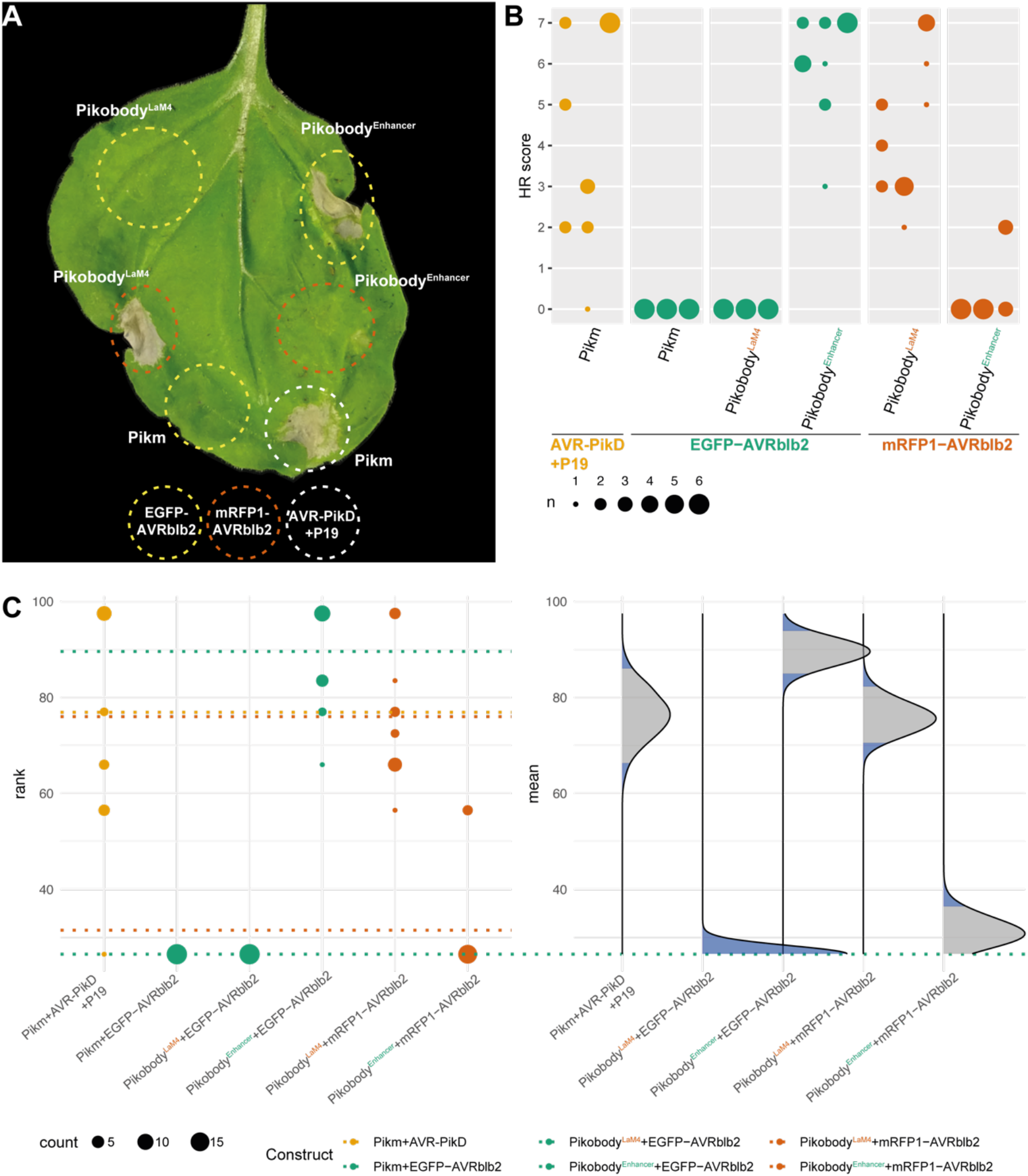
Pikobodies specifically trigger HR in response to EGFP- or mRFP1-tagged *Phytophtora infestans* effector AVRblb2. (**A**) Pikobody^Enhancer^ and Pikobody^LaM4^ specifically recognize EGFP- or mRFP1-tagged AVRblb2, respectively. Representative *N. benthamiana* leaf infiltrated with constructs indicated in white and infiltration sites circled with a dashed line, photographed 4 days after infiltration. Yellow, Orange or white color show sites co-infiltrated with EGFP-AVRblb2, mRFP1-AVRblb2 or AVR-PikD+P19, respectively. (**B**) HR scores visualized as dots plot, where the size of a dot is proportional to the number of samples with the same score (n) within the same replicate (1 to 3). The experiment was repeated three times with six internal replicates; each column represents a combination of constructs (labelled on the bottom). (**C**) Statistical analysis of hypersensitive response shown in (**B**) conducted using an estimation method using besthr R library (MacLean, 2020). The wild-type Pikm pair with EGFP-tagged AVRblb2 was used as a negative control, while the wild-type Pikm pair with the cognate effector AVR-PikD and the silencing inhibitor p19 was used as a positive control for HR. The left panel represents the ranked data (dots) and their corresponding mean (dashed line), with the size of a dot proportional to the number of observations with that specific value. The panel on the right shows the distribution of 1000 bootstrap sample rank means, with the blue areas illustrating the 0.025 and 0.975 percentiles of the distribution. The difference is considered significant if the ranked mean for a given condition falls within or beyond the blue percentile of the mean distribution for another condition.

**Fig. S5.**
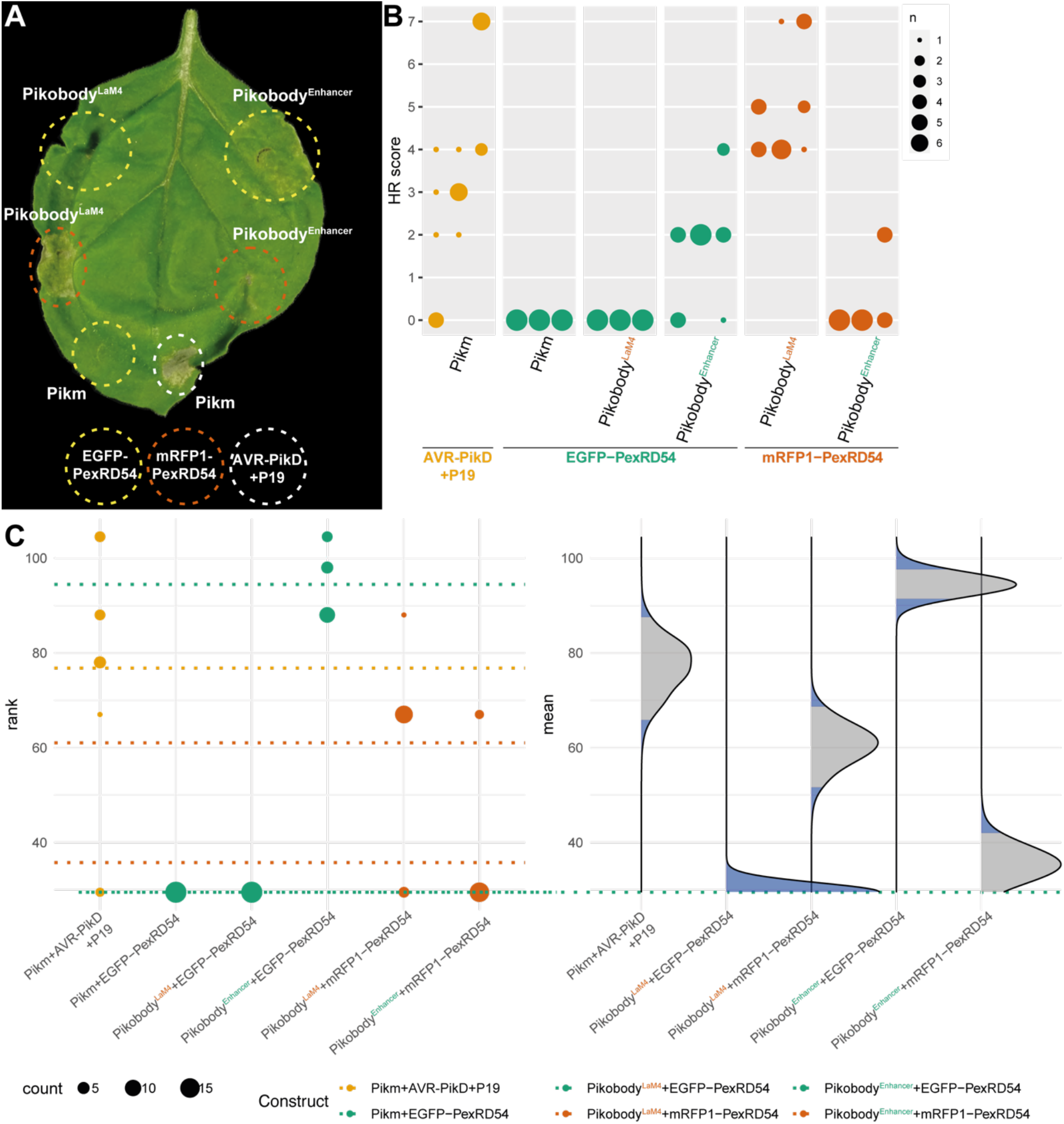
Pikobodies specifically trigger HR in response to EGFP- and mRFP1-tagged *P. infestans* effector PexRD54. (**A**) Pikobody^Enhancer^ and Pikobody^LaM-4^ specifically recognize EGFP- or mRFP1-tagged PexRD54, respectively. Representative *N. benthamiana* leaf infiltrated with constructs indicated in white and infiltration sites circled with a white dashed line, photographed 4 days after infiltration. Yellow, Orange or white color show sites co-infiltrated with EGFP-PexRD54, mRFP1-PexRD54 or AVR-PikD+P19, respectively. (**B**) HR scores visualized as dots plot, where the size of a dot is proportional to the number of samples with the same score (n) within the same replicate (1 to 3). The experiment was repeated three times with six internal replicates; each column represents a combination of constructs (labelled on the bottom). (**C**) Statistical analysis of hypersensitive response shown in (**B**) conducted using an estimation method using besthr R library (MacLean, 2020). The wild-type Pikm pair with EGFP-tagged PexRD54 was used as a negative control, while the wild-type Pikm pair with the cognate effector AVR-PikD and the silencing inhibitor p19 was used as a positive control for HR. The left panel represents the ranked data (dots) and their corresponding mean (dashed line), with the size of a dot proportional to the number of observations with that specific value. The panel on the right shows the distribution of 1000 bootstrap sample rank means, with the blue areas illustrating the 0.025 and 0.975 percentiles of the distribution. The difference is considered significant if the ranked mean for a given condition falls within or beyond the blue percentile of the mean distribution for another condition.

**Fig. S6.**
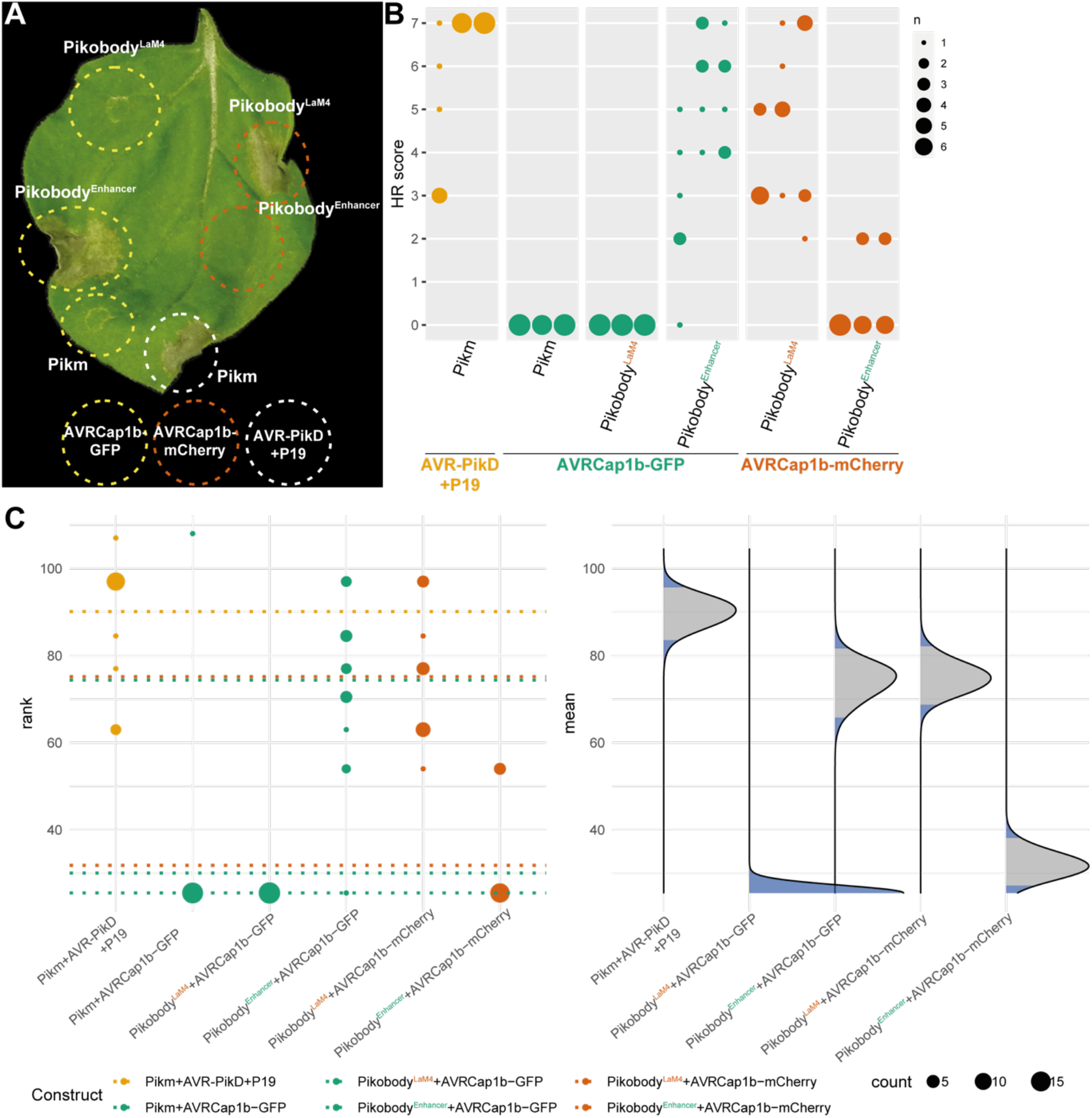
Pikobodies specifically trigger HR in response to GFP- and mCherry-tagged *P. infestans* effector AVRCap1b. (**A**) Pikobody^Enhancer^ and Pikobody^LaM4^ specifically recognize GFP- or mCherry-tagged AVRcap1b, respectively. Representative *N. benthamiana* leaf infiltrated with constructs indicated in white and infiltration sites circled with a white dashed line, photographed 4 days after infiltration. Yellow, Orange or white color show sites co-infiltrated with AVRCap1b-GFP, AVRCap1b-mCherry- or AVR-PikD+P19, respectively. (**B**) HR scores visualized as dots plot, where the size of a dot is proportional to the number of samples with the same score (n) within the same replicate (1 to 3). The experiment was repeated three times with six internal replicates; each column represents a combination of constructs (labelled on the bottom). (**C**) Statistical analysis of hypersensitive response shown in (**B**) conducted using an estimation method using besthr R library (MacLean, 2020). The wild-type Pikm pair with GFP-tagged AVRCap1b was used as a negative control, while the wild-type Pikm pair with the cognate effector AVR-PikD and the silencing inhibitor p19 was used as a positive control for HR. The left panel represents the ranked data (dots) and their corresponding mean (dashed line), with the size of a dot proportional to the number of observations with that specific value. The panel on the right shows the distribution of 1000 bootstrap sample rank means, with the blue areas illustrating the 0.025 and 0.975 percentiles of the distribution. The difference is considered significant if the ranked mean for a given condition falls within or beyond the blue percentile of the mean distribution for another condition.

**Fig. S7.**
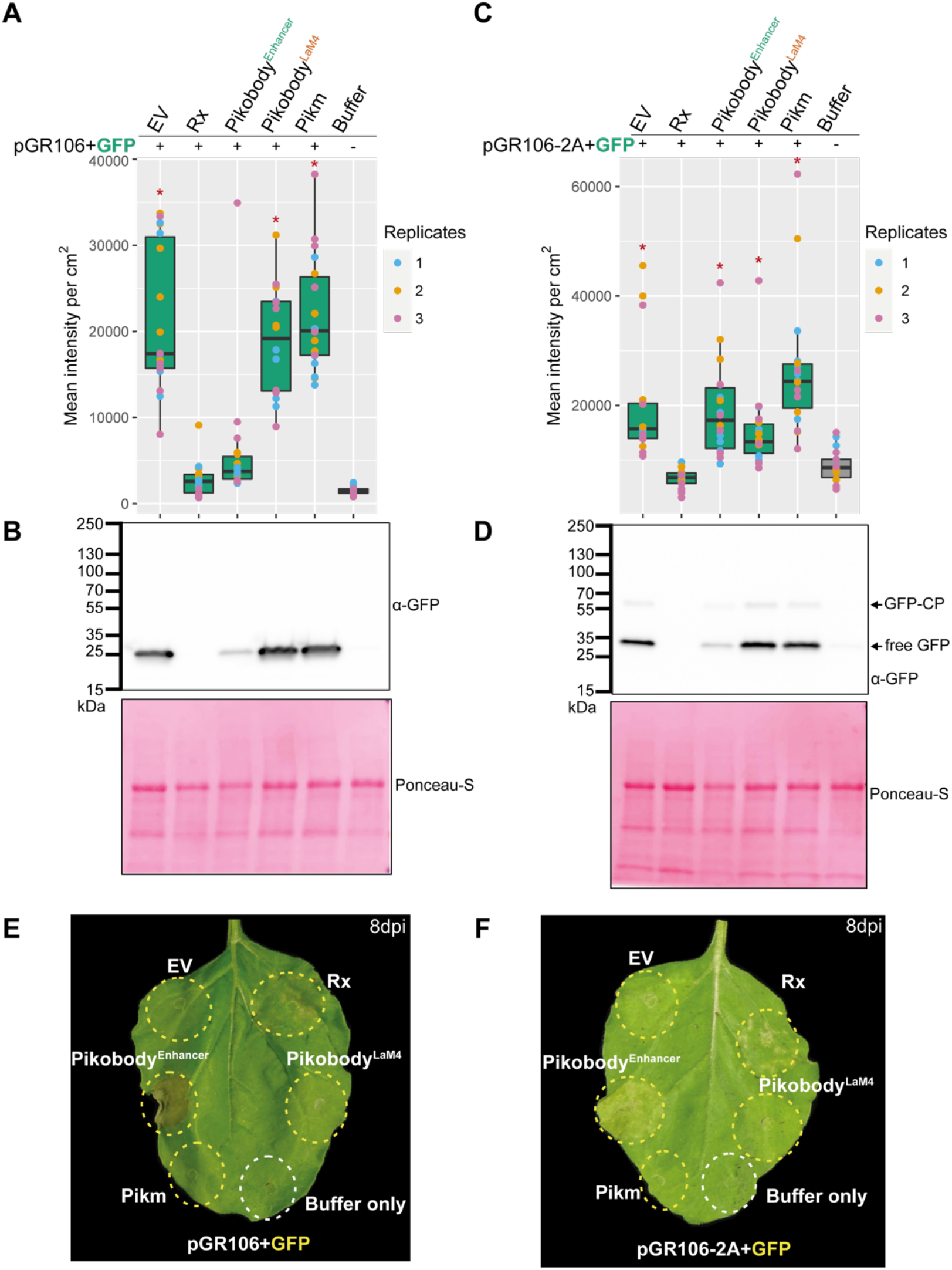
Pikobody^Enhancer^ confers resistance against two other engineered variants of Potato Virus X expressing GFP. (**A**) Specific reduction in fluorescence intensity of PVX expressed GFP in the presence of Pikobody^Enhancer^. GFP mean fluorescence intensity per cm^2^ measured in *N. benthamiana* leaves 4 days post-infiltration was used as a proxy for PVX viral load in each infiltration site for PVX engineered to express GFP from between the triple gene block and the coat protein (pGR106-GFP). The boxplot summarizes data obtained in three independent experiments (replicates) with six infiltration site per construct and per experiment. Red asterisks show significant differences between Buffer only (no PVX added, negative control) and tested constructs in the presence of pGR106-GFP (Dunnett’s test, pvalue < 0.001). Empty vector was used as a positive control for viral load and Buffer only as a negative control for viral load. We also used Rx, a resistance gene recognizing the coat protein from PVX as a positive control for PVX resistance. (**B**) Specific reduction of PVX expressed GFP accumulation in the presence of Pikobody^Enhancer^. GFP accumulation was used as proxy to evaluate viral load from pGR106-GFP. For the immunoblot analysis, total protein was extracted 2 days after inoculation with pGR106-GFP in the presence of the tested constructs. Empty vector was used as a positive control for viral load and Buffer only as a negative control. GFP was detected using anti-GFP antibody. Ponceau S staining shows equal protein loading across samples. (**C**) No reduction in fluorescence intensity of PVX expressed GFP fused to the coat protein in the presence of Pikobody^Enhancer^. GFP mean fluorescence intensity per cm^2^ measured in *N. benthamiana* leaves 4 days post-infiltration was used as a proxy for PVX viral load in each infiltration site for PVX engineered to express GFP fused to the coat protein using the porcine teschovirus-1 2A “self-cleaving” peptide (pGR106-2A-GFP) leading to approximately 50 % of free GFP and 50 % of GFP fused to the viral coat protein (CP). Boxplots and analysis as in (**A**). (**D**) Specific reduction of PVX expressed GFP accumulation fused to the coat protein in the presence of Pikobody^Enhancer^. GFP accumulation was used as proxy to evaluate viral load from pGR106-2A-GFP. Immunoblot analysis as in (**B**) with black arrows pointing to GFP fused to coat protein (GFP-CP) and free GFP. (**E-F**) Specific HR triggered by the two version of PVX expressing GFP in the presence of Pikobody^Enhancer^. Representative *N. benthamiana* leaves infiltrated with appropriate constructs photographed 8 days after infiltration. The infiltration site for each construct is labelled on the picture and circled with a yellow or white dashed line for pGR106+GFP (**E**) and pGR106-2A+GFP (**F**) or no PVX, respectively.

**Fig. S8.**
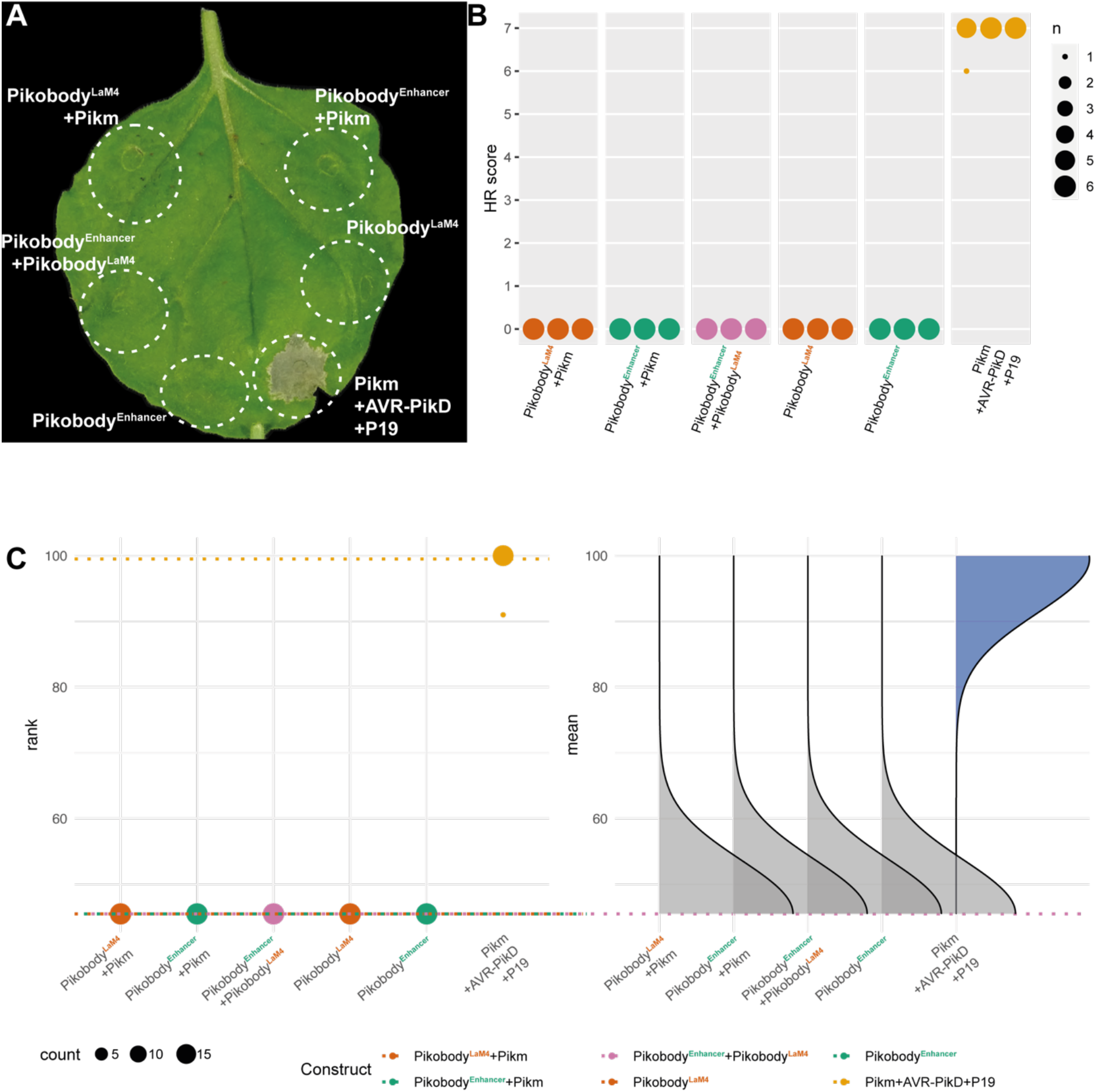
Co-expressed Pikobodies are not autoactive. (**A**) Stacking of Pikobodies does not result in autoimmunity. Representative *N. benthamiana* leaf infiltrated with constructs indicated in white and infiltration sites circled with a white dashed line, photographed 4 days after infiltration. (**B**) HR scores visualized as dots plot, where the size of a dot is proportional to the number of samples with the same score (n) within the same replicate (1 to 3). The experiment was repeated three times with six internal replicates; each column represents a combination of constructs (labelled on the bottom). Wild-type Pikm pair co-expressed with cognate effector AVR-PikD and silencer suppressor p19 was used as a positive control and Pikobody^Enhancer^ or Pikobody^LaM4^ expressed on their own as negative controls (Figure S1). (**C**) Statistical analysis of hypersensitive response shown in (**B**) conducted using an estimation method using besthr R library (MacLean, 2020). Pikobody^LaM4^ was used as a negative control (Figure S1), while the wild-type Pikm pair with the cognate effector AVR-PikD and the silencing inhibitor p19 was used as a positive control for HR. The left panels represent the ranked data (dots) and their corresponding mean (dashed line), with the size of a dot proportional to the number of observations with that specific value. The panels on the right show the distribution of 1000 bootstrap sample rank means, with the blue areas illustrating the 0.025 and 0.975 percentiles of the distribution. The difference is considered significant if the ranked mean for a given condition falls within or beyond the blue percentile of the mean distribution for another condition.

**Fig. S9.**
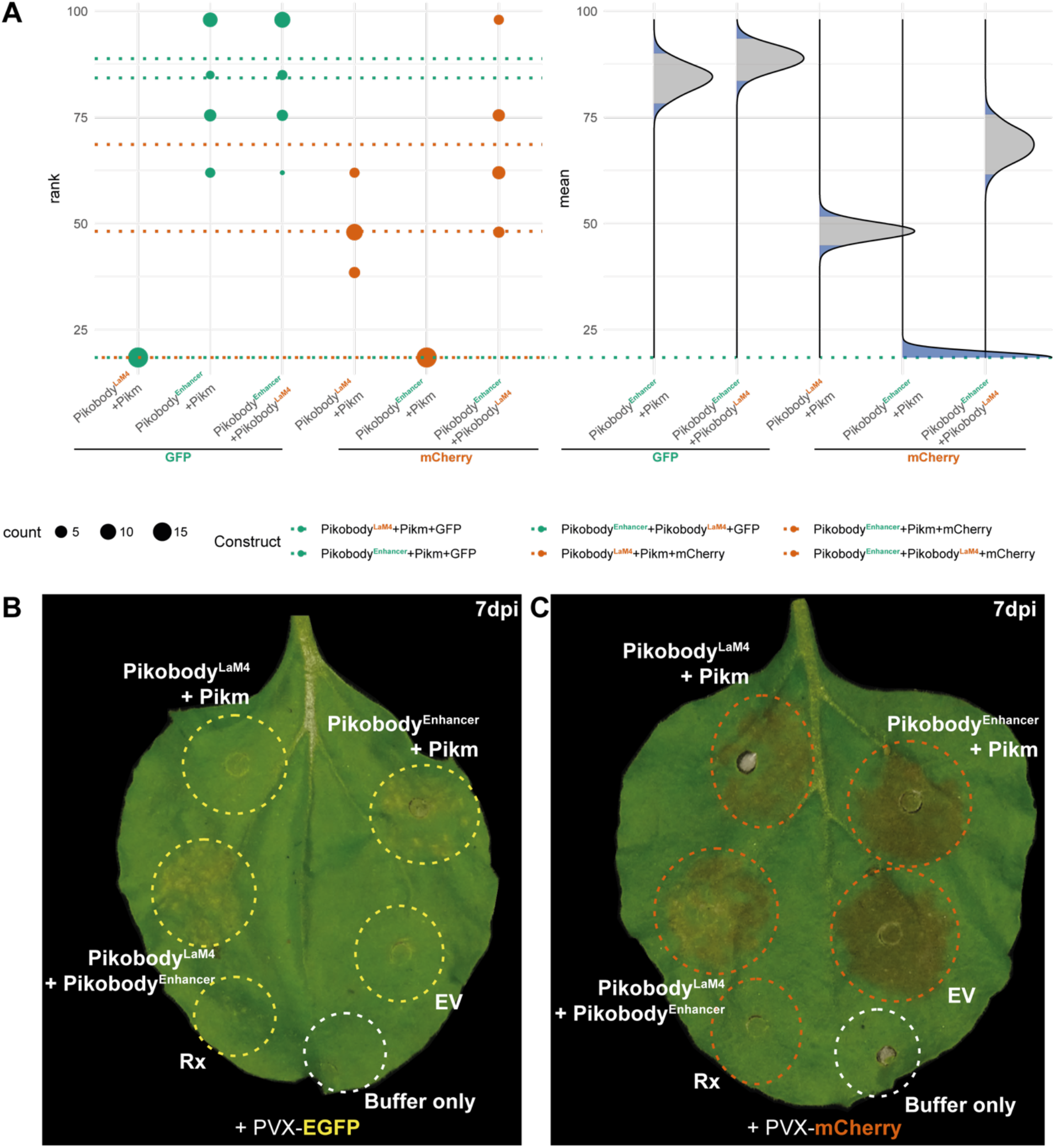
Statistical analysis of Pikobody stacking and HR phenotype. (**A**) Statistical analysis of hypersensitive response shown in **Fig. 3** conducted using an estimation method using besthr R library (MacLean, 2020). The wild-type Pikm pair with Pikobody^LaM4^ and GFP was used as a negative control, while the wild-type Pikm pair with the cognate effector AVR-PikD and the silencing inhibitor p19 was used as a positive control for HR. The left panel represents the ranked data (dots) and their corresponding mean (dashed line), with the size of a dot proportional to the number of observations with that specific value. The panel on the right shows the distribution of 1000 bootstrap sample rank means, with the blue areas illustrating the 0.025 and 0.975 percentiles of the distribution. The difference is considered significant if the ranked mean for a given condition falls within or beyond the blue percentile of the mean distribution for another condition. (**B-C**) Specific HR triggered by PVX expressed EGFP (GFP, B) and mCherry (C) in the presence of stacked Pikobody^Enhancer^ and Pikobody^LaM4^, respectively. Representative *N. benthamiana* leaves infiltrated with appropriate constructs photographed 7 days after infiltration. The infiltration site for each construct is labelled on the picture and all sites but “Buffer only” were co-infiltrated with PVX-GFP (left) or PVX-mCherry (right). Yellow, orange abd white colors depict the sites infiltrated with PVX-GFP, PVX-mCherry and no PVX, respectively. The purple color on the infiltration sites on (C) is due to a high expression of mCherry.

**Fig. S10.**
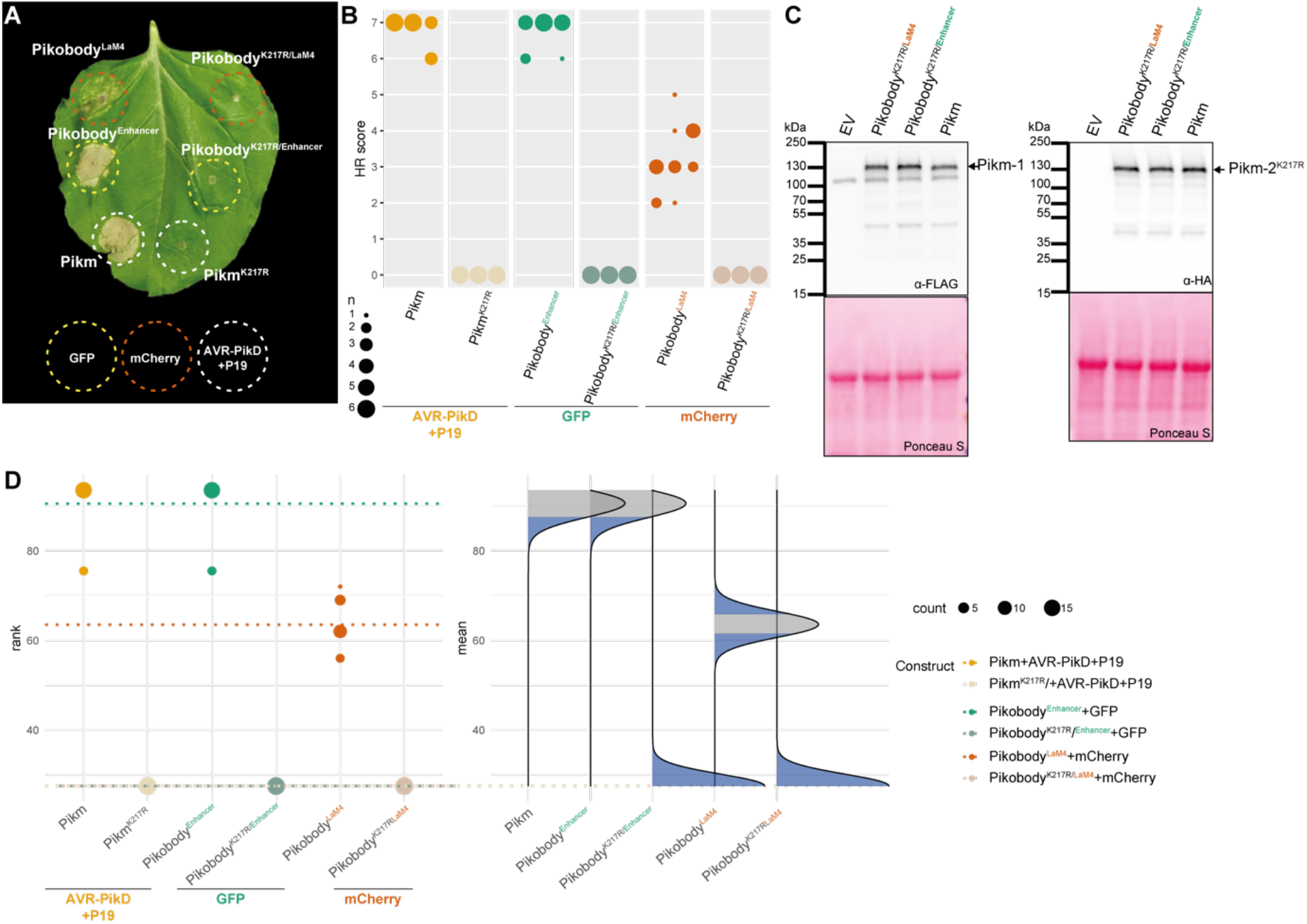
Pikobody-mediated HR requires the ploop motif. (**A**) An intact Pikm-2 P-loop motif is required for Pikobody-mediated HR. Representative *N. benthamiana* leaf infiltrated with constructs indicated in white and infiltration sites circled with a dashed line, photographed 4 days after infiltration. Yellow, Orange or white color shows sites co-infiltrated with GFP, mCherry or AVR-PikD+P19, respectively. Wild-type Pikm pair co-expressed with cognate effector AVR-PikD and silencer suppressor p19 was used as a positive control and Pikm^K217R^ P-loop mutant (mutation in Pikm-2) co-expressed with cognate effector AVR-PikD and silencer suppressor p19 was used as a negative control. (**B**) HR scores visualized as dots plot, where the size of a dot is proportional to the number of samples with the same score (n) within the same replicate (1 to 3). The experiment was repeated three times with six internal replicates; each column represents a combination of constructs (labelled on the bottom). The statistical analyses of these results are presented in (**D**). (**C**) Pikm-2^K217R^ P-loop mutant protein accumulates. For the immunoblot analysis, total protein was extracted 2 days after transient expression of Pikobody^Enhancer^, Pikobody^LaM4^ and the wild-type Pikm pair. Pikm-1 and the Pikm-1 Enhancer and LaM-4 were C-terminally 6xHis-3xFLAG-tagged and detected using anti-FLAG antibody, while Pikm-2^K217R^ was C-terminally 6xHA-tagged and detected using anti-HA antibody. Empty vector was used as a negative control. Back arrows point to the band corresponding to Pikm-1 (anti-FLAG blot) and Pikm-2^K217R^ (anti-HA blot). (**D**) Statistical analysis of hypersensitive response showed in (**B**) conducted using an estimation method using besthr R library (MacLean, 2020). The Pikm pair with the Pikm-2^K217R^ P-loop mutant was used as as a negative control, while the wild-type Pikm pair with the cognate effector AVR-PikD and the silencing inhibitor p19 was used as a positive control for HR. The left panel represents the ranked data (dots) and their corresponding mean (dashed line), with the size of a dot proportional to the number of observations with that specific value. The panel on the right shows the distribution of 1000 bootstrap sample rank means, with the blue areas illustrating the 0.025 and 0.975 percentiles of the distribution. The difference is considered significant if the ranked mean for a given condition falls within or beyond the blue percentile of the mean distribution for another condition.

**Table S1: Description of constructs and *Agrobacterium* strains**.

Details of the constructs used in this study with associated selection antibiotic, purpose, publication, agrobacterium strain and OD_600_ used for infiltrations.

**Table S2: Hypersensitive cell death quantification**.

HR scores reported by construct, experiment replicate (Experiment# in the table) and associated figure. We described HR scoring in the corresponding section of the Materials and Methods.

**Table S3: Fluorescence intensity quantification**.

Fluorescence intensity recorded for each infiltration site with the ImageQuant LAS 4000 luminescent imager (GE Healthcare Life Sciences) as described in the corresponding Materials and Methods section. Each line represents a construct co-infiltrated with a given PVX strain or no PVX (buffer only) with corresponding fluorescence per cm^2^ calculated from the fluorescent intensity measured in each infiltration site. We indicated the different experiment replicates in the column “Experiment#” and the corresponding Figures in the last two columns.

